# Insights into the mechanism of oligodendrocyte protection and remyelination enhancement by the integrated stress response

**DOI:** 10.1101/2023.01.23.525156

**Authors:** Yanan Chen, Songhua Quan, Vaibhav Patil, Rejani B. Kunjamma, Haley M. Tokars, Eric D. Leisten, Jonah Chan, Yvette Wong, Brian Popko

## Abstract

CNS inflammation triggers activation of the integrated stress response (ISR). We previously reported that prolonging the ISR protects remyelinating oligodendrocytes and promotes remyelination in the presence of inflammation (Chen et al., *eLife*, 2021). However, the exact mechanisms through which this occurs remain unknown. Here, we investigated whether the ISR modulator Sephin1 in combination with the oligodendrocyte differentiation enhancing reagent bazedoxifene (BZA) is able to accelerate remyelination under inflammation, and the underlying mechanisms mediating this pathway. We find that the combined treatment of Sephin1 and BZA is sufficient to accelerate early-stage remyelination in mice with ectopic IFN-γ expression in the CNS. IFN-γ, which is a critical inflammatory cytokine in multiple sclerosis (MS), inhibits oligodendrocyte precursor cell (OPC) differentiation in culture and triggers a mild ISR. Mechanistically, we further show that BZA promotes OPC differentiation in the presence of IFN-γ, while Sephin1 enhances the IFN-γ-induced ISR by reducing protein synthesis and increasing RNA stress granule formation in differentiating oligodendrocytes. Finally, the ISR suppressor 2BAct is able to partially lessen the beneficial effect of Sephin1 on disease progression, in an MS mouse model of experimental autoimmune encephalitis (EAE). Overall, our findings uncover distinct mechanisms of action of BZA and Sephin1 on oligodendrocyte lineage cells under inflammatory stress, suggesting that a combination therapy may effectively promote restoring neuronal function in MS patients.

## INTRODUCTION

Multiple sclerosis (MS) is an immune-mediated demyelinating disease that causes neurological impairment and frequently results in significant disability (Compston and Coles, 2002; Frohman et al., 2006; Reich et al., 2018). Although the etiology of MS is still unknown, the consensus is that central nervous system (CNS) inflammation is a major driver of MS pathology. It is well accepted that activated autoreactive myelin-specific T cells migrate into the CNS, are reactivated locally, and release cytokines leading to oligodendrocyte and myelin damage (Lassmann, 2018; Titus et al., 2020).

Oligodendrocytes produce a vast amount of myelin membrane and thus are particularly sensitive to cellular homeostatic changes that result from diverse stresses, including CNS inflammation (Clayton and Popko, 2016; Lin and Popko, 2009; Volpi et al., 2016). To cope with inflammatory stress, an innate cytoprotective response called the integrated stress response (ISR) is triggered in oligodendrocytes, centered by the phosphorylation of the eukaryotic translation initiation factor 2 alpha (eIF2α) (Pakos-Zebrucka et al., 2016; Way and Popko, 2016; Way et al., 2015). Phosphorylated eIF2α (p-eIF2α) inhibits the activity of eukaryotic translation initiation factor 2B (eIF2B), and diminishes global translation by preventing ternary complex formation, thus reducing the protein load on the endoplasmic reticulum (ER). Concurrently, p-eIF2α selectively upregulates translation of chaperones and protective proteins, including the activating transcription factor 4 (ATF4), ATF5, C/EBP homologous protein (CHOP), and protein growth arrest and DNA-damage inducible (GADD34), aiding cellular survival and recovery (Zhang and Kaufman, 2008). GADD34 is recruited to protein phosphatase 1 (PP1), which dephosphorylates p-eIF2α ensuring translation reinitiation. Sephin1, a recently-identified small molecule, is able to prolong the ISR by inhibiting GADD34-PP1c complex activity, thereby prolonging the phosphorylation of eIF2α (Crespillo-Casado et al., 2017; Das et al., 2015). We previously showed that Sephin1 protects oligodendrocytes against inflammatory stress in a mouse model of MS, experimental autoimmune encephalomyelitis (EAE) (Chen et al., 2019). However, the exact mechanism by which Sephin1 protects oligodendrocytes from inflammation remains unknown.

Remyelination is known to be incomplete in MS as remyelinating oligodendrocytes, both pre-existing and newly formed, are challenged by an inflammatory environment (Ruffini et al., 2004). Inflammation diminishes the capacity of oligodendrocyte precursor cells (OPCs) to develop into mature oligodendrocytes and remyelinate demyelinated lesions (Franklin and Ffrench-Constant, 2008; Starost et al., 2020). The T cell cytokine interferon-gamma (IFN-γ) is considered to be a critical inflammatory mediator of MS pathogenesis (Patel and Balabanov, 2012). Using *GFAP-tTA; TRE-IFN-γ* transgenic mice in which IFN-γ is secreted in a doxycycline (dox)-dependent manner from astrocytes, we have previously shown that IFN-γ expression in the CNS suppresses remyelination following cuprizone-induced oligodendrocyte toxicity. However, prolonging the ISR through Sephin1 treatment or mutating GADD34 protects remyelinating oligodendrocytes and increases the level of remyelination (Chen et al., 2021). In the same inflammatory demyelination/remyelination model, we have also shown that combining Sephin1 with the oligodendrocyte differentiation enhancing reagent bazedoxifene (BZA) further increases myelin thickness, but not the number of remyelinating oligodendrocytes or remyelinated axons (Chen et al., 2021). Nevertheless, whether the combined treatment is able to accelerate remyelination under inflammatory conditions remains untested and the underlying mechanisms of the observed effects have not been fully elucidated.

Here, we investigated the remyelination effect of Sephin1 and BZA at an early stage of remyelination using the *GFAP-tTA; TRE-IFN-γ* transgenic mouse model along with cuprizone-induced demyelination. We demonstrate that the combined treatment of Sephin1 and BZA show an additive effect on promoting early-stage remyelination in the presence of inflammation. Using primary OPC cultures exposed to IFN-γ, we further show that BZA promotes OPC differentiation while Sephin1 boosts the IFN-γ-induced ISR by reducing protein synthesis and increasing RNA stress granule formation in the differentiating cells. In addition, we provide evidence using an EAE mouse model that 2BAct, a highly selective eIF2B activator and an ISR suppressor, partially blunts the beneficial effect of the ISR enhancer Sephin1 on disease progression. Collectively, our results illustrate that Sephin1’s mechanism is distinct from that of BZA to protect oligodendrocytes against inflammation by inhibiting translation initiation and activating stress granule formation.

## RESULTS

### Combined treatment of Sephin1 and BZA accelerates early-stage remyelination in the presence of IFN-γ

In our previous study, using a *GFAP-tTA;TRE-IFN-γ* double-transgenic mouse model of cuprizone demyelination/remyelination, we did not observe additional benefits from the combined treatment of Sephin1 and BZA on the number of oligodendrocytes and remyelinated axons present at three weeks after cuprizone withdrawal (Chen et al., 2021). However, given that remyelination in the treatment groups had already reached pre-lesion levels after three weeks of remyelination, we reasoned that examining earlier recovery time-points might reveal differences in the treatment groups. Thus, to examine whether the combined treatment confers any benefit at an earlier stage of remyelination, we utilized the same *GFAP-tTA;TRE-IFN-γ* double-transgenic mouse model along with cuprizone exposure (Chen et al., 2021; Lin et al., 2006). We removed the doxycycline (Dox) chow to induce CNS expression of IFN-γ and placed the mice on a diet of 0.2% cuprizone chow to trigger demyelination at six weeks of age (Figure 1a). Sephin1 (8 mg/kg, i.p.) and/or BZA (10 mg/kg, gavage) were administered daily to the mice beginning three weeks after starting the cuprizone diet, which represents a point of peak demyelination. After five weeks of cuprizone exposure, the mice were placed back on a normal diet for two weeks (Figure 1a). Only mice with the highest level of IFN-γ expression in the CNS were selected for further examination at two weeks after cuprizone withdrawal (Figure 1-figure supplement 1). Using electron microscopy, we observed that only the combined Sephin1/BZA group presented a significantly higher number of remyelinated axons (*p<0.05) than the vehicle group in the corpus callosum of the IFN-γ expressing mice at the early-remyelination stage (Figure 1b,c). This is in contrast to our previous finding at the late-stages of remyelination, 3 weeks after cuprizone withdrawal, during which both individual and combined treatments displayed more remyelinated axons than the vehicle group (Chen et al., 2021). Moreover, the g-ratios of myelinated axons were significantly lower in the combination treatment group than the vehicle-treated group (*p<0.05) and single treatment groups (**p<0.01 vs Sephin1, *p<0.05 vs BZA) (Figure 1b,c), indicating that axons of the combination treated mice had thicker myelin.

**Figure 1.**
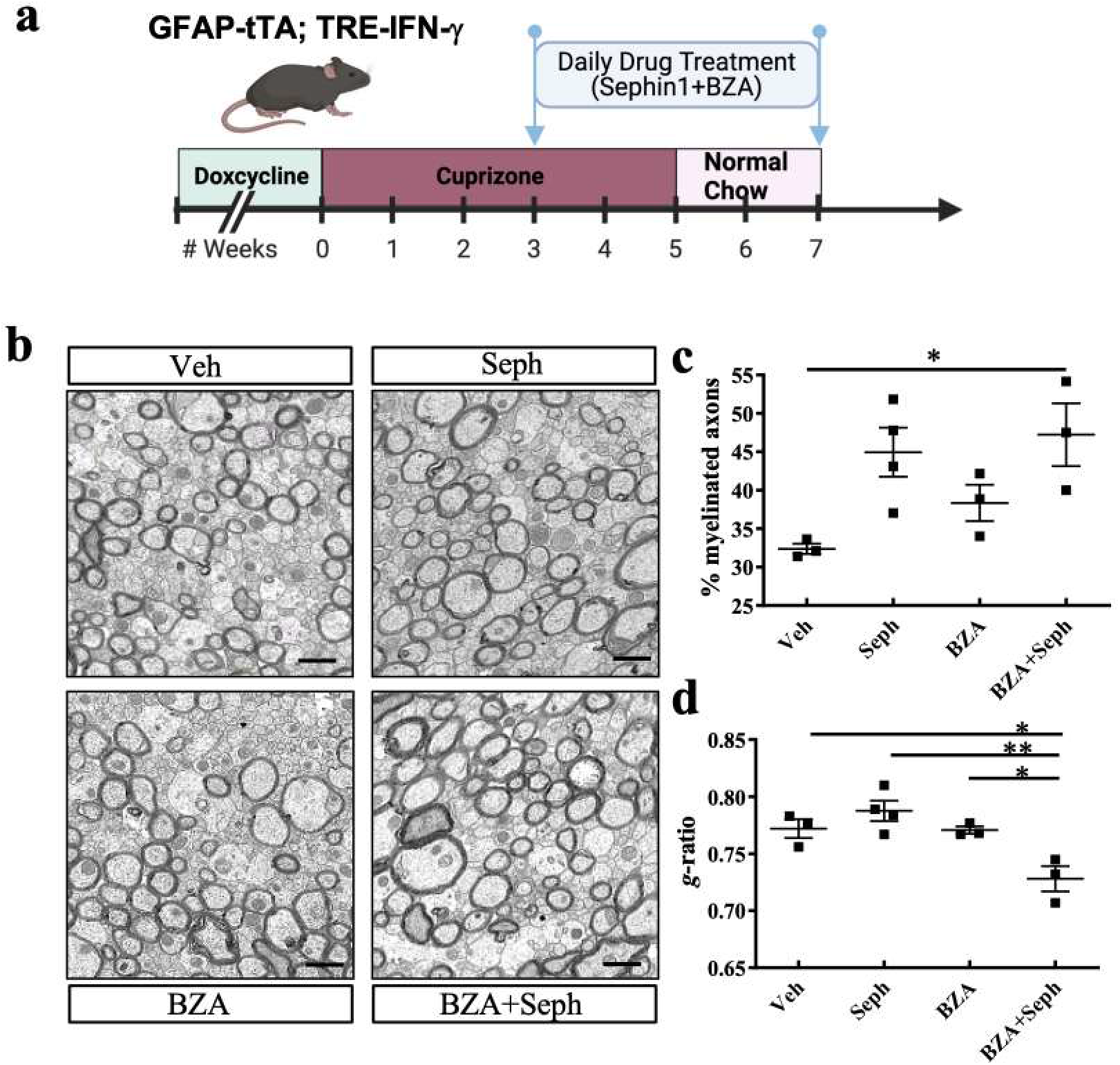
Combined treatment of Sephin1 and BZA promotes early–stage remyelination in the presence of IFN-γ. (**a**) Cuprizone demyelination/remyelination model of GFAP-tTA;TRE-IFN-γ with designed treatment. Drug treatment was started after three weeks of cuprizone exposure and lasted to the early remyelination stage. (**b**) Corpora callosa of GFAP-tTA;TRE-IFN-γ were taken for EM processing at two weeks after cuprizone withdrawal. Representative EM image of the corpus callosum. Scale bar: 1μm. Quantifications of percentage of remyelinated axons (**c**) and g-ratios of axons (**d**) in the corpus callosum areas in the presence of IFN-γ. *p < 0.05, significance based on ANOVA. Data represent an average of 3–4 mice per group (mean ± SEM).

We also examined the status of remyelinating oligodendrocytes after two weeks. Using immunofluorescent staining, we found that mice receiving combined Sephin1/BZA and individual BZA treatment had more CC1+ mature oligodendrocytes than vehicle treatment (*p<0.05) in the corpus callosum of the IFN-γ expressing mice (Figure 2a,b). Interestingly, the intensity of phosphorylated eIF2α (p-eIF2α) was significantly increased in CC1+ oligodendrocytes when treated with Sephin1 or Sephin1/BZA rather than with vehicle, suggesting that Sephin1 enhanced the ISR in remyelinating oligodendrocytes in the presence of IFN-γ (Figure 2a,c). Together, these findings indicate that the combined treatment of Sephin1 and BZA drives the remyelination process in the presence of IFN-γ faster than either treatment alone.

**Figure 2.**
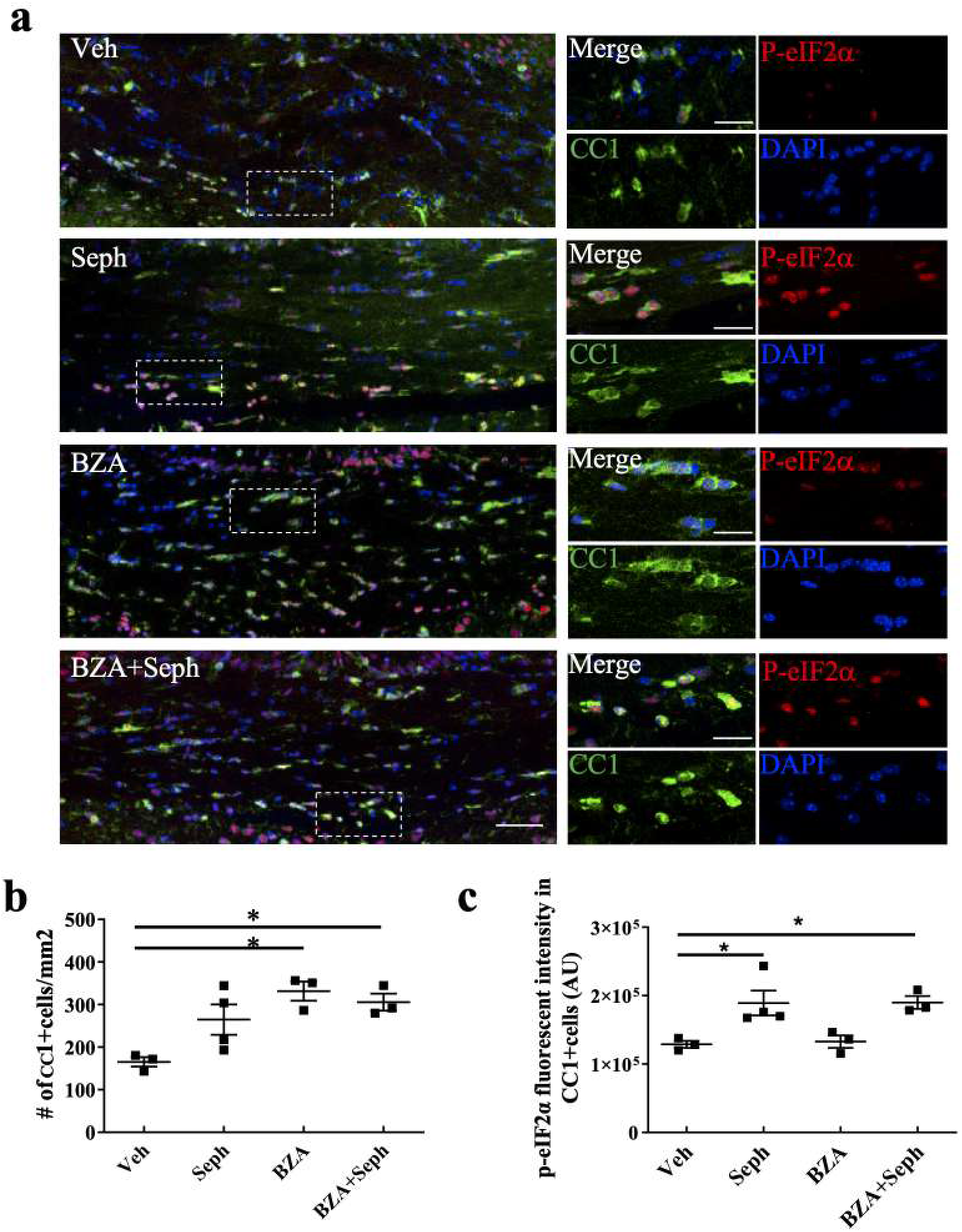
Combined treatment of Sephin1 and BZA increases the number of remyelinating oligodendrocytes in the early–stage remyelination. (**a**) Immunostaining of the corpus callosum with anti-cc1, anti-p-eIF2α and DAPI. Scale bar: 50μm. Higher magnification images are taken from the boxed areas. Scale bar = 20 μm. (**b**) Quantification of cells positive for CC1 in the corpus callosum. (**c**) Quantification of the fluorescent intensity of p-eIF2α staining in the CC1+ oligodendrocytes. *p < 0.05, significance based on ANOVA. Data represent an average of 3–4 mice per group (mean ± SEM).

### Combined treatment of Sephin1 and BZA does not affect OPC proliferation but promotes OPC differentiation in the presence of IFN-γ

Remyelination is a regenerative process that requires the recruitment and proliferation of oligodendrocyte progenitor cells (OPCs) to demyelinated areas. This is followed by their differentiation into mature remyelinating oligodendrocytes to form functional myelin sheaths (Franklin and Ffrench-Constant, 2008). To determine whether the combined treatment of Sephin1 and BZA promotes OPC proliferation, we examined the corpus callosum of the IFN-γ expressing mice with the OPC marker PDGFRα and the proliferation marker Ki67 at two weeks of remyelination. Combined or individual treatments of Sephin1 and BZA had no effect on the number of total OPCs or proliferative OPCs (PDGFRα+ and Ki67+) in the demyelinated lesions (Figure 3a,b). Next, we interrogated the influence of Sephin1 and BZA on differentiation of OPCs isolated from postnatal mouse pups with or without the exposure of IFN-γ *in vitro*. Significantly fewer OPCs differentiated into MBP+ oligodendrocytes in the cultures exposed to IFN-γ (+IFN-γ; NT) than non-IFN-γ treated cultures (-IFN-γ; NT) (#p<0.05, unpaired t test) after 24 hours (Figure 3c,d). Interestingly, when treated with Sephin1 (+IFN-γ; Seph), OPC differentiation reached the control level of non-IFN-γ (-IFN-γ; NT) cultures (Figure 3c,d). These data suggest that OPC differentiation is inhibited by IFN-γ, while Sephin1 permits OPC differentiation in the presence of IFN-γ. Combined Sephin1/BZA and BZA alone, however, promoted OPCs to differentiate into MBP+ oligodendrocytes both in the absence and presence of IFN-γ (*p<0.05 vs BZA, **p<0.01 vs Sephin1/BZA, Figure 3c,d). No additional difference was observed between the combination Sephin1/BZA treatment and individual BZA treatment. Similar results were found after 48 hours and 72 hours of treatment (Figure 3-figure supplement 2). Overall, this data shows that Sephin1 permits OPC differentiation in the presence of inflammatory stress, while BZA promotes OPC differentiation in control or inflammatory conditions.

**Figure 3.**
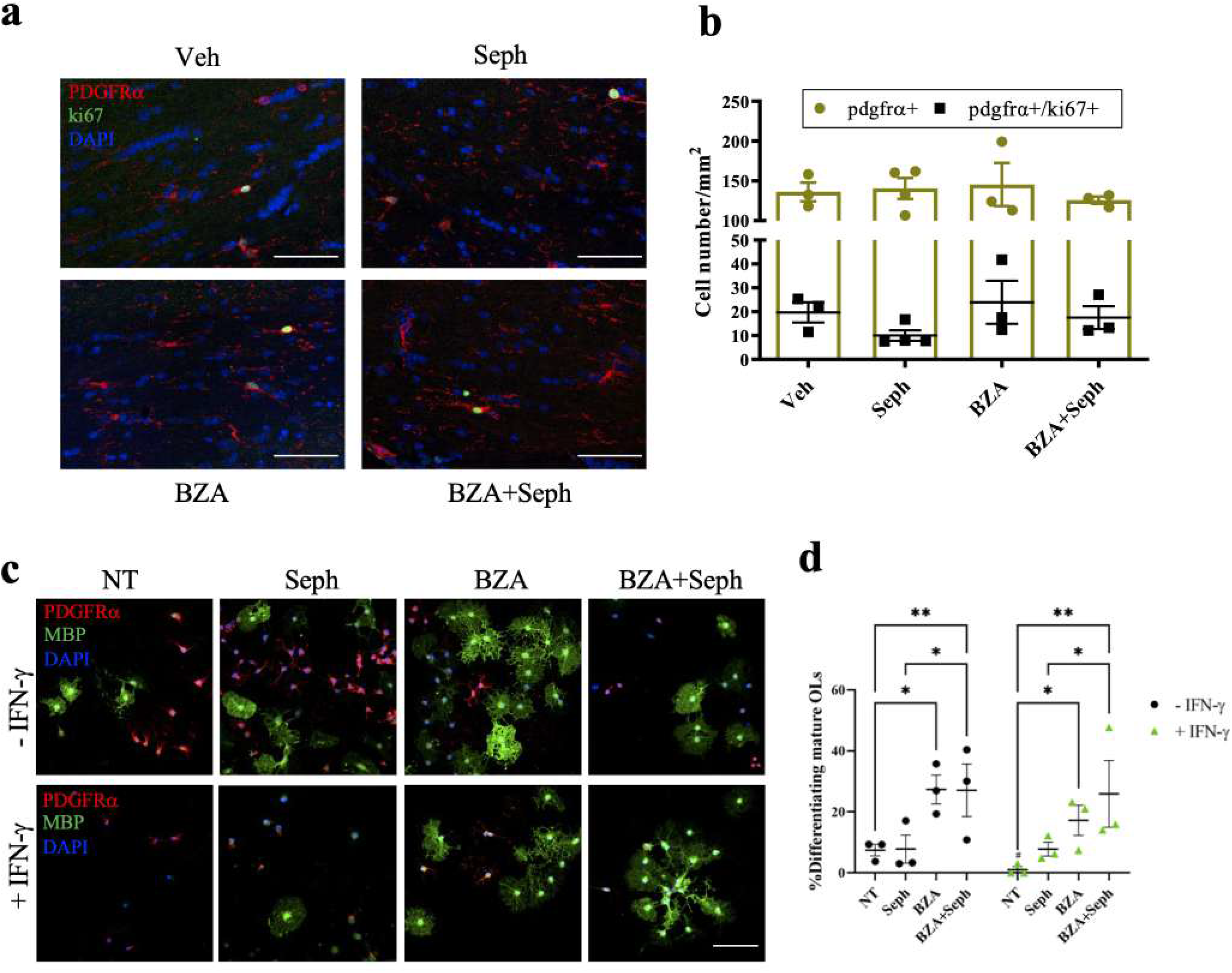
Combined treatment of Sephin1 and BZA does not change OPC proliferation but promotes OPC differentiation. (**a**) Corpora callosa of GFAP-tTA;TRE-IFN-γ were taken for immunostaining at two weeks after cuprizone withdrawal. Immunostaining of corpus callosum with anti-PDGFR-α, anti-Ki67 and DAPI. Scale bar: 50μm. (**b**) Quantification of cells positive for PDGFR-α and cells positive with both PDGFR-α and Ki67 at the corpus callosum. (**c**) PDGFR-α and MBP immunostaining of OPCs in cultures that were seeded for 24 hours. Cells exposed to IFN-γ (+ IFN-γ) were treated with either nontreatment (NT), Sephin1 (Seph), BZA, or BZA plus Seph1. Cells were not exposed to IFN-γ (-IFN-γ) as controls. Scale bar: 100μm. (**d**) Quantification of percentage of cells positive for MBP (differentiating oligodendrocytes) over the total number of oligodendrocyte lineage cells. *p < 0.05, **p < 0.01.

### eIF2α phosphorylation induction by IFN-γ is not sufficient to inhibit protein synthesis in oligodendrocytes

Before exploring the influence of Sephin1 on stressed oligodendrocytes, we first needed to better understand the inflammatory environment surrounding oligodendrocytes and other CNS resident cells. T-cell secreted IFN-γ is considered to be a critical inflammatory mediator in MS (Patel and Balabanov, 2012). Both our *in vivo* and *in vitro* models incorporated IFN-γ to mimic the inflammatory environment of MS. Previously, our studies have shown that IFN-γ activates the ISR in differentiating OPCs as indicated by increased p-eIF2α levels (Chen et al., 2019; Lin et al., 2005). However, the downstream molecular mechanisms by which IFN-γ influences oligodendrocytes remain unclear.

First, we exposed purified differentiating OPCs to IFN-γ or a well-known ER stressor thapsigargin (TG) and compared the expression of downstream targets of the ISR by Western-blot. TG induces ER stress by inhibiting Ca^2+^-ATPase and thereby disturbing ER calcium homeostasis (Lytton et al., 1991; Sehgal et al., 2017). Levels of p-eIF2α significantly increased rapidly with the TG treatment, peaking at one hour (***p<0.0001) (Figure 4a,c). Similarly, IFN-γ increased p-eIF2α expression but at a much slower rate, starting at eight hours (***p<0.0001) and peaking at 20 hours (p<0.0001) (Figure 4b,d). We also examined downstream stress-induced ISR signaling components, which aid in cell survival and recovery. ATF4, GADD34, CHOP, and BIP levels in oligodendrocytes significantly increased after two hours of TG treatment, indicating a fully activated ISR (Figure 4e,g). The levels were comparable to those observed in positive control NIH 3T3 cells treated with TG. Surprisingly, IFN-γ treatment did not alter the expression of these ISR components at any of the time points examined (Figure 4f,h).

**Figure 4.**
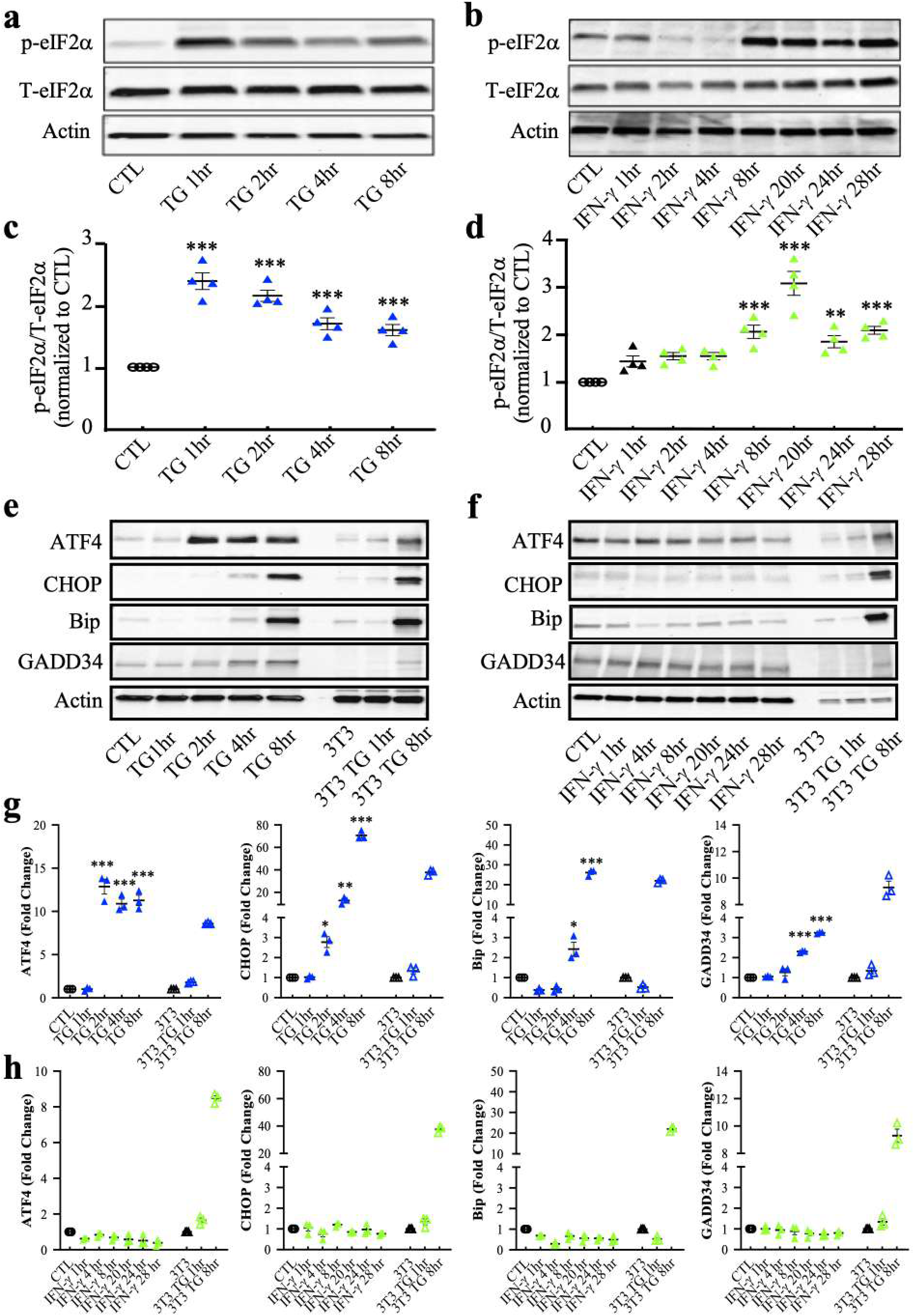
Upregulation of downstream ISR proteins in thapsigargin-treated oligodendrocytes, but not in IFN-γ treated oligodendrocytes. **(a)** Immunoblotting analysis of p-eIF2α and total eIF2α in thapsigargin (TG) treated developing oligodendrocytes. **(b)** Immunoblotting analysis of p-eIF2α and total eIF2α in IFN-γ treated developing oligodendrocytes. **(c** and **d)** Quantification of p-eIF2α levels with respect to Western blots **a** and **b**. **(e)** Western blot analysis of the ISR response (ATF4, BIP, GADD34, and CHOP) in TG treated developing oligodendrocytes. **(f)** Western blot analysis of the ISR response in IFN-γ treated developing oligodendrocytes. As a positive control, NIH 3T3 cells were also incubated with TG for 1 or 8 hrs. (**g** and **h**) Quantification of Western blots **e** and **f.** Data are mean ± SEM. *p < 0.05, **p < 0.01, ***p < 0.001. **Figure 4-source data 1**. Western-blots images of downstream ISR proteins in oligodendrocytes exposed to thapsigargin or IFN-γ.

Phosphorylation of eIF2α mediates translational control, causing an overall reduction in translation initiation. To examine the effect of p-eIF2α caused by IFN-γ on protein synthesis, we utilized a puromycin assay based on the structural resemblance between puromycin and the 3’ end of aminoacylated tRNA (aa-tRNA). We added puromycin to cultured OPCs, where puromycin instead of aa-tRNA entered the ribosome’s A-site and accepted the nascent peptide chain, leading to translation termination and the release of nascent chains bearing a 3’ puromycin. These puromycilated peptides were detected by Western-blots using anti-puromycin antibodies, generating a laddering pattern that reflects newly synthesized proteins (Schmidt et al., 2009). Following the pattern of p-eIF2α levels, TG significantly reduced puromycin incorporated protein synthesis at one hour (***p<0.001) and eight hours of treatment (***p<0.001) (Figure 5a,c). In contrast, we did not observe any changes in protein synthesis by IFN-γ treatment (Figure 5b,d). In addition, to determine whether IFN-γ interferes with ISR activation caused by ER stress, we also treated cells with a combination of IFN-γ and TG. We found that TG was still capable of inducing eIF2α phosphorylation and reducing protein synthesis in the presence of IFN-γ (***p<0.001) (Figure 5-figure supplement 3). Overall, these data indicate that although the ISR is activated in oligodendrocytes in response to the presence of IFN-γ, a full ISR response is not achieved.

**Figure 5.**
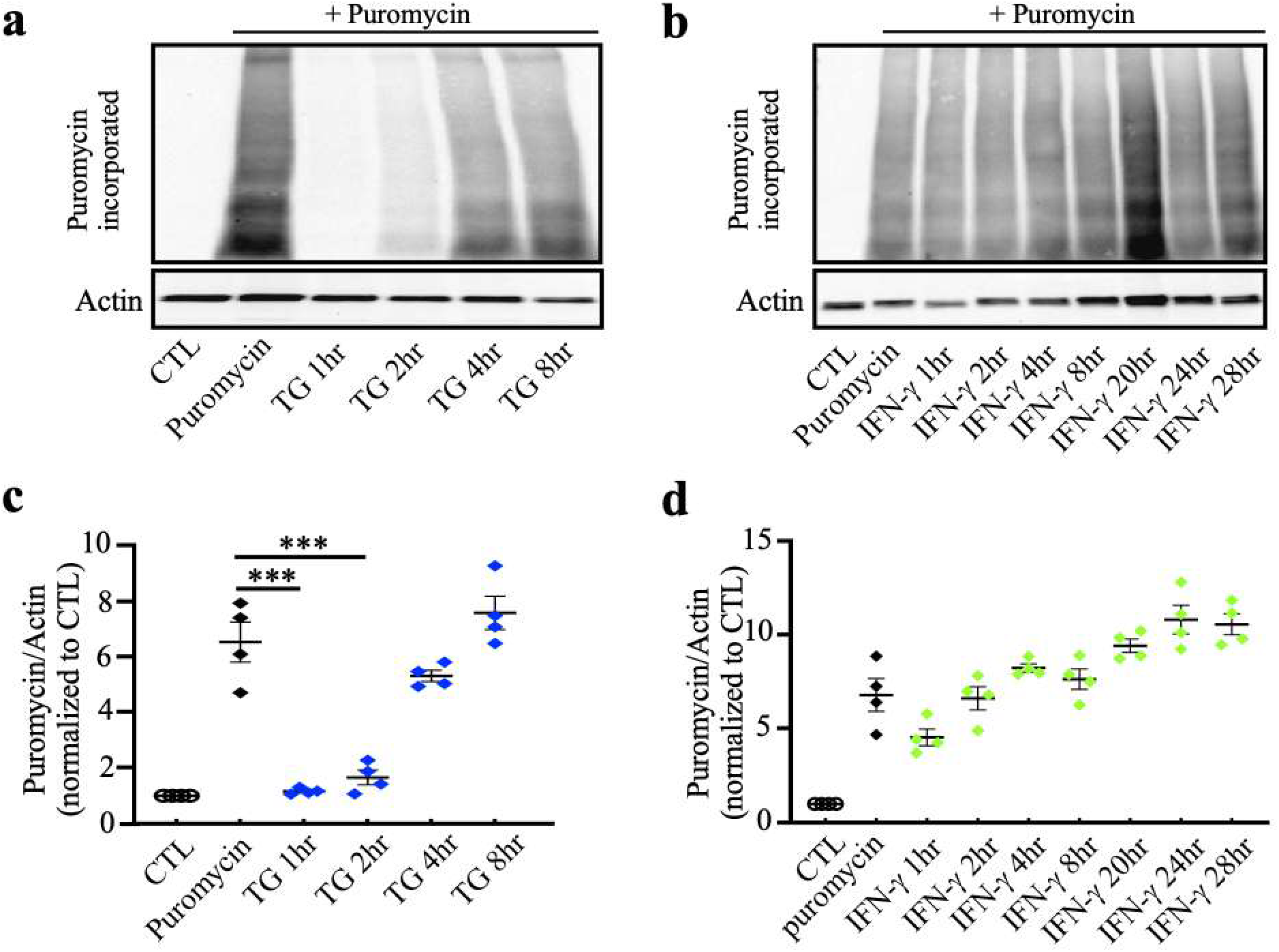
Inhibition of overall cellular protein translation in thapsigargin treated oligodendrocytes, but not in IFN-γ treated oligodendrocytes. **(a)** Western blots showing elevated p-eIF2α levels in thapsigargin (TG) treated oligodendrocytes. **(b)** Western blots showing elevated p-eIF2α levels in IFN-γ treated oligodendrocytes. **(c** and **d)** Quantification of Western blots **a** and **b**. **(e)** Incorporation of puromycin into TG was detected using an anti-puromycin antibody. **(f)** Incorporation of puromycin into IFN-γ was detected using an anti-puromycin antibody. Treatment of puromycin alone served as a positive control. (**g** and **h**) Quantification of Western blots **e** and **f.** Data are mean ± SEM. *p < 0.05, **p < 0.01, ***p < 0.001. **Figure 5-source data 1**. Western-blots images of p-eIF2α and puromycin in oligodendrocytes exposed to thapsigargin or IFN-γ.

### Sephin1 induces translation reduction in IFN-γ stressed oligodendrocytes by upregulating the phosphorylation of eIF2α and the formation of RNA stress granules

Previously, we showed that Sephin1 prolongs eIF2α phosphorylation in IFN-γ-stressed oligodendrocytes *in vivo* (Chen et al., 2019). Here, we also observed increased p-eIF2α in remyelinating oligodendrocytes of IFN-γ expressing mice treated with Sephin1 (Figure 2a,c). This prolonged phosphorylation of p-eIF2α during inflammatory stress via Sephen1 may provide protection to oligodendrocytes, but the downstream molecular mechanism is unclear. We therefore examined p-eIF2α levels and protein synthesis in IFN-γ-exposed OPCs treated with Sephin1 by immunoblotting. Sephin1 further increased the phosphorylation of eIF2α in IFN-γ stressed oligodendrocytes at eight hours and significantly at 20 hours (*p<0.05) (Figure 6a,b). Strikingly, Sephin1 also significantly lessened protein synthesis in these cells at 20 hours (*p<0.05) (Figure 6c,d). In addition, Sephin1 reduced protein synthesis while enhancing the level of p-eIF2α in TG-stressed oligodendrocytes (Figure 6). A full-blown ISR reduces the levels of overall protein synthesis, while activating translation of a set of mRNA encoding for transcription factors and other cytoprotective proteins. Given that p-eIF2α levels were increased by Sephin1, we examined the expression levels of downstream components of the ISR. Although there was not a significant increase in ATF4 expression, the levels of GADD34 were increased after 24 hours of Sephin1 treatment in IFN-γ stressed oligodendrocytes (Figure 6-figure supplement 4).

**Figure 6.**
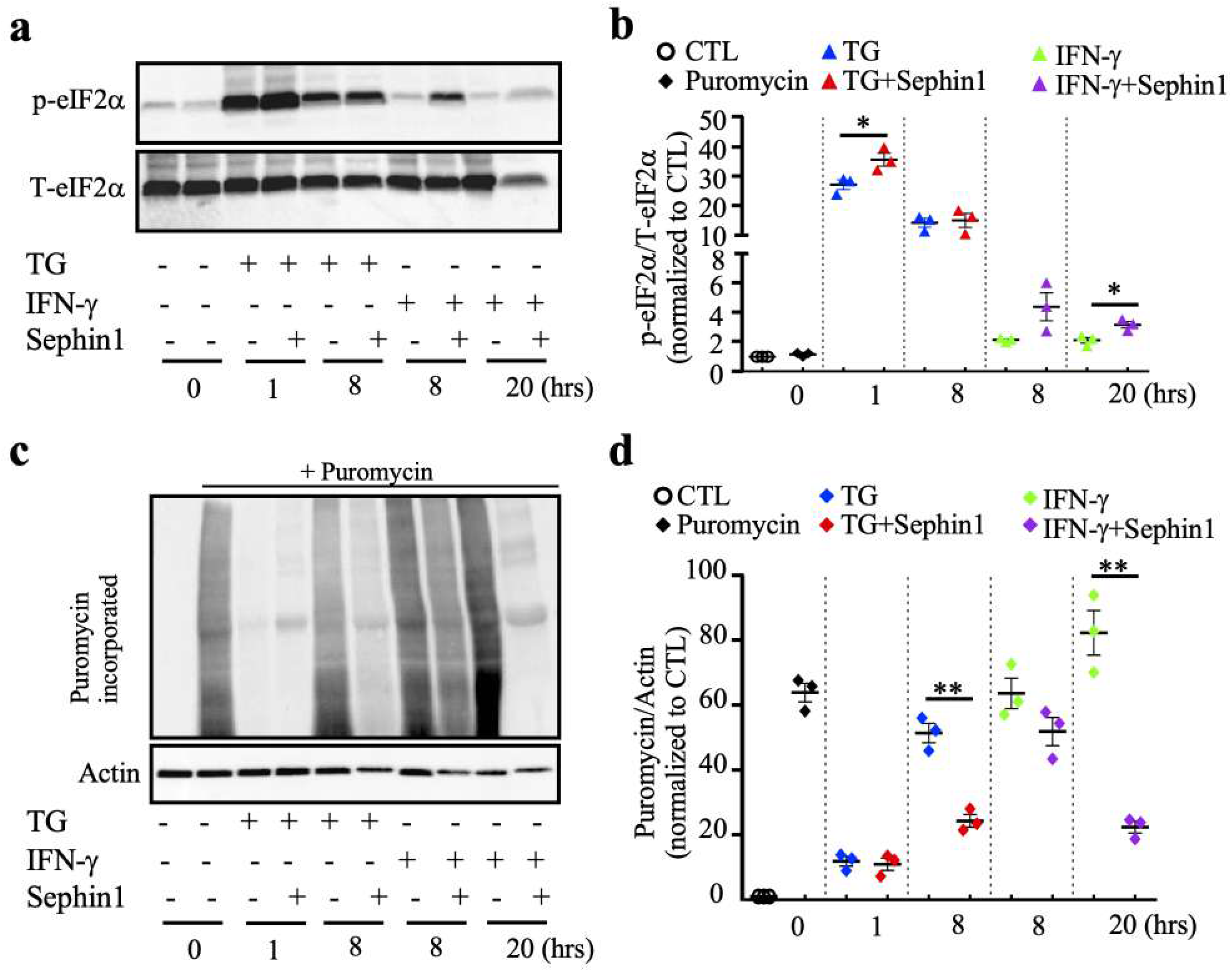
Sephin1 prolongs the ISR response and reduces the overall protein translation in mouse oligodendrocytes. **(a)** Western blot of oligodendrocytes treated either by thapsigargin (TG) or IFN-γ with or without Sephin1. **(b)** Quantification of Western blots for p-eIF2α levels. **(c)** Western blot of puromycin incorporation assay using an anti-puromycin antibody. OPCs were differentiated in differentiation media overnight and then treated with either TG or IFN-γ with or without Sephin1. Puromycin was added in the last 30 min before harvesting for puromycin labeling. **(d)** Quantification of Western blot. Data are mean ± SEM from three biological isolations and technique replicates. *p < 0.05, **p < 0.01, ***p < 0.001. **Figure 6-source data 1**. Western-blots images of p-eIF2α and puromycin in oligodendrocytes exposed to IFN-γ with or without Sephin1 treatment.

Translation inhibition by phosphorylation of eIF2α generates ribosome-free mRNA that becomes sequestered into RNA stress granules (Glauninger et al., 2022; Hofmann et al., 2021; Kedersha et al., 1999). To identify if stress granule assembly occurs in oligodendrocytes treated with TG or IFN-γ + Sephin1, we stained treated oligodendrocytes at different timepoints with the RasGAP SH3-domain-binding protein 1 (G3BP1), a key component for stress granule formation (Tourrière et al., 2003). G3BP1 was present diffusively in the cytoplasm of cultured oligodendrocytes without treatment (CTL), but clearly formed granules were present in sodium arsenite as well as TG treated cells after one hour (Figure 7a). In contrast, G3BP1 granules were not found in cells that were exposed for one or eight hours to IFN-γ (Figure 7a, b). Remarkably, distinct G3BP1 positive granules appeared in oligodendrocytes exposed to both IFN-γ and Sephin1 after eight hours (Figure 7b). Stress granules were dispersed by 20 or 48 hours of exposure to the dual treatment (Figure 7 – figure supplement 5a). To further confirm the presence of stress granules, we assessed the G3BP1 stress granules by superresolution microscopy (Structured Illumination Microscopy, SIM). Here too, we observed G3BP1 granules only in cells treated with both IFN-γ and Sephin1 at the 8- and 16-hour timepoints, and not in the single treatment cells (Figure 7c; Figure 7-figure supplement 5b). Based on the observation above, we infer that Sephin1 promotes oligodendrocytes survival under inflammatory attack by reducing protein synthesis and/or promoting stress granules assembly.

**Figure 7.**
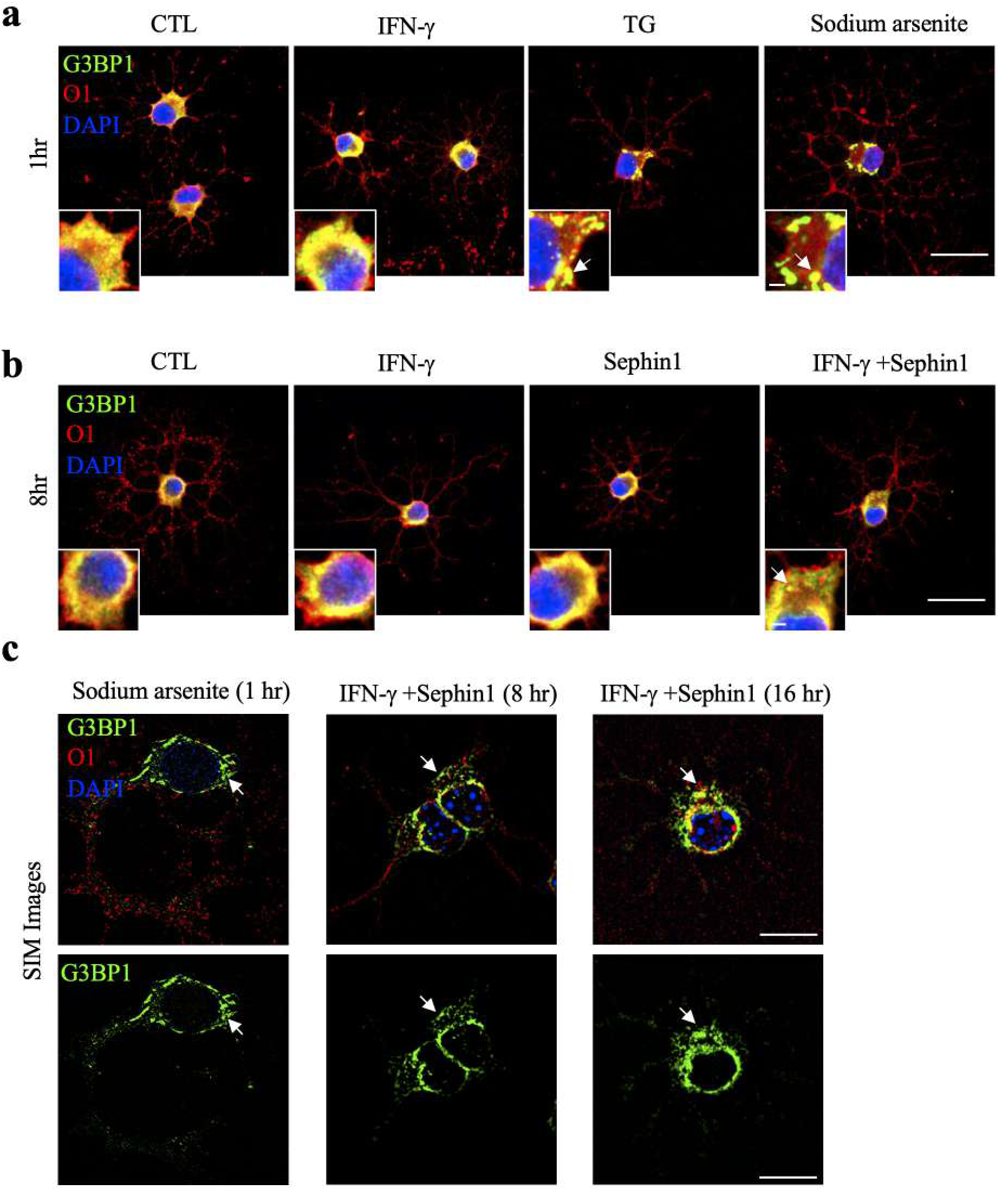
Stress granule formation in IFN-γ stressed oligodendrocytes when treated with Sephin1. (**a-b**) Representative confocal images of the stress granule marker G3BP1 (green) and oligodendrocyte marker O1 (red) in purified oligodendrocytes. Purified OPCs were incubated in differentiation media overnight and then were either untreated (CTL) or treated 1hr with IFN-γ, TG or sodium arsenite for one hour (**a**) and 8 hours (**b**). Scale bars indicate 20 μm and 2μm (inserts). White arrows point to G3BP1 puncta. (**c**) Representative SIM images of G3BP1 (green) and O1 (red). Nuclei were stained with DAPI (blue). Scale bar indicates 10 μm. White arrows point to G3BP1 puncta.

### 2BAct partially counteracts the beneficial effect of Sephin1 on EAE

Phosphorylated eIF2α acts as a competitive inhibitor of eIF2B, which slows protein synthesis (Krishnamoorthy et al., 2001). A highly selective eIF2B activator, 2BAct, has been shown to blunt the ISR by antagonizing the inhibitory effect of p-eIF2α on eIF2B (Wong et al., 2019). We therefore hypothesized that if Sephin1 protects oligodendrocytes from inflammation by prolonging ISR-induced elevated levels of p-eIF2α, this effect could be inhibited by 2BAct. To test this *in vivo*, we conducted daily Sephin1 treatments on EAE mice provided with normal chow or chow containing 2BAct. As previously reported (Chen et al., 2019), Sephin1 delayed the onset of EAE peak symptoms (Figure 8). In contrast, Sephin1 treated mice fed chow containing 2BAct developed EAE symptoms significantly faster than those on normal chow (at PID 15 and PID 16; p<0.05) (Figure 8), although the beneficial impact of Sephin1 was not completely abrogated by 2BAct. These findings further attest that Sephin1 provides CNS protection against inflammation by enhancing the ISR.

**Figure 8.**
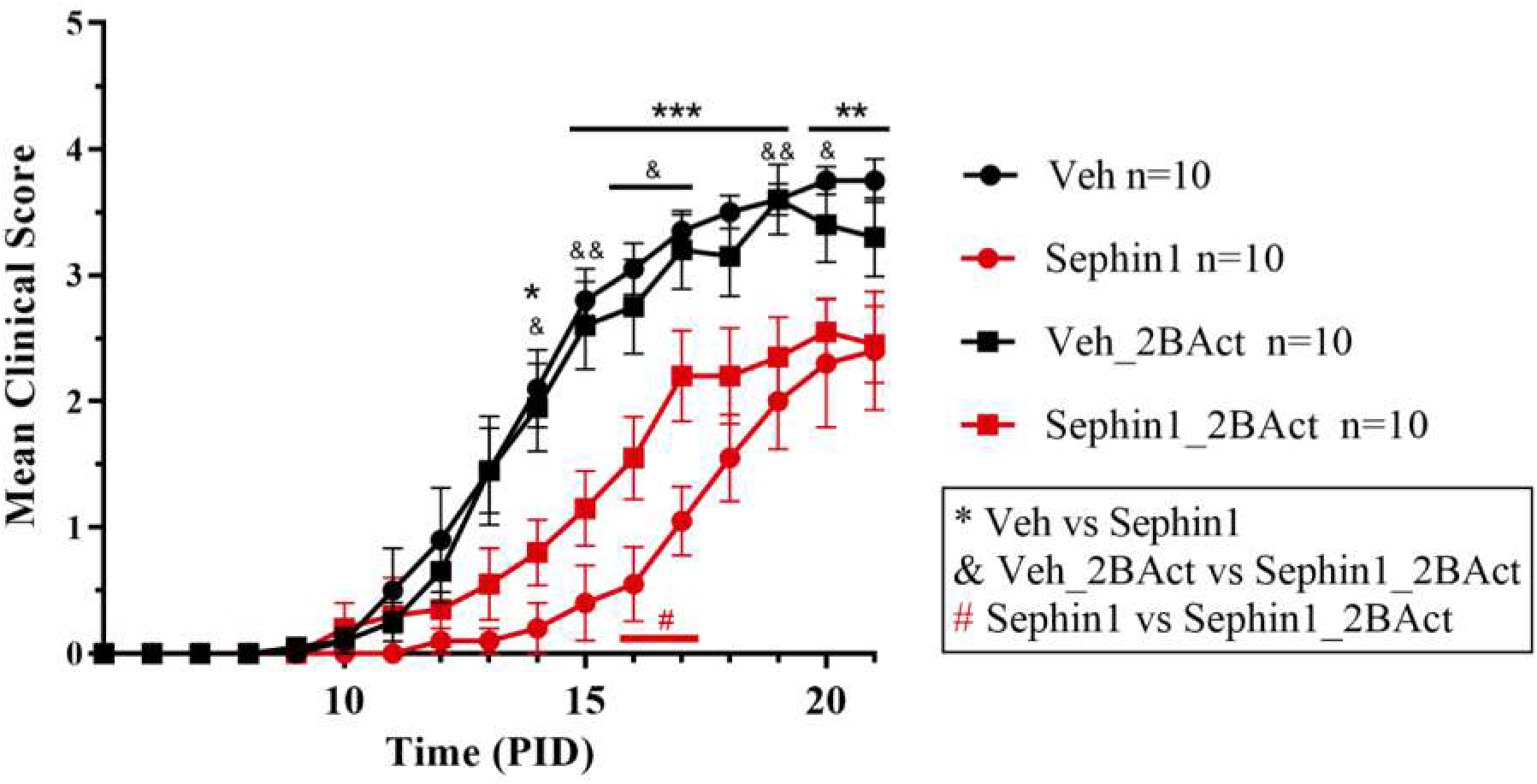
2BAct partially blocks the protective effect of Sephin1 on EAE. Mean clinical scores of C57BL/6J female mice immunized with MOG35-55/CFA to induce chronic EAE. Mice were either on normal chow or chow containing 2BAct, treated with vehicle or 8 mg/kg Sephin1. n=10 in each group. *p<0.05, **p<0.01, ***p<0.001; &p<0.05, &&p<0.01. #p<0.05.

## DISCUSSION

In this study, we have shown that the combination of Sephin1 and BZA accelerates the remyelination process faster than Sephin1 or BZA individually in the presence of an adverse environment created by the inflammatory cytokine IFN-γ. Our results indicate that this additive effect is derived from Sephin1 and BZA’s different mechanisms of action on the remyelinating oligodendrocyte lineage cells. We show that BZA promotes OPC differentiation in the presence of IFN-γ, whereas Sephin1 permits oligodendrocyte development in these conditions. Moreover, our results indicate that Sephin1 extends the level of IFN-γ triggered ISR by enhancing p-eIF2α levels, which leads to reduced overall protein synthesis and the formation RNA stress granules. In addition, we show that an ISR suppressor 2BAct, which reverses the effects of p-eIF2α, can partially block the protective effects of Sephin1 on EAE, indicating that Sephin1 safeguards oligodendrocytes against inflammation by enhancing the ISR. Our observation that the combined treatment of Sephin1 and BZA resulted in more rapid and robust remyelination in the presence of inflammation is critical because remyelination provides protection against axonal degeneration secondary to demyelination. More rapid remyelination is likely to preserve more axons.

Remyelination restores neuronal function and prevents axonal degeneration, but remyelination is either incomplete or absent in MS (Fancy et al., 2010; Franklin and Goldman, 2015). Efforts to promote remyelination have largely focused on promoting OPC differentiation into remyelinating oligodendrocytes (Franklin and Simons, 2022; Franklin et al., 2021). Nevertheless, stimulating repair is challenging in the hostile environment created by CNS inflammation, during which OPC differentiation efficacy largely declines (Franklin and Ffrench-Constant, 2017). BZA, a third-generation selective estrogen receptor modulator, has been found to enhance OPC differentiation in response to focal demyelination, independent of it estrogen receptor (Rankin et al., 2019). In our recent study, we found that BZA promotes remyelination in an inflammatory remyelination model as well (Chen et al., 2021). Here, we further showed that BZA can promote OPC differentiation in an inflammatory milieu and that remyelination is further accelerated by the ISR modulator Sephin1.

In our inflammatory demyelination/remyelination model, we utilized IFN-γ as the inflammatory stressor. However, the mechanism by which IFN-γ leads to oligodendrocyte abnormalities remains poorly understood. It has been well documented that the T-cell-derived pleiotropic cytokine IFN-γ plays a critical, albeit complex, role in the development and progression of MS and EAE (Goverman, 2009; Lees and Cross, 2007; Lin et al., 2007; Popko et al., 1997). We previously reported that the inhibitory effects of IFN-γ on the remyelination process are associated with an activated stress response in oligodendrocytes, although the mechanisms are unclear (Lin et al., 2006). ISR induction not only reduces translation in general but also promotes translation of a set of cytoprotective proteins (Holcik and Sonenberg, 2005; Zhang and Kaufman, 2008). Here, we observed that despite significant eIF2α phosphorylation in cells exposed to IFN-γ, there was no inhibition of global protein synthesis or an increase in the level of translation of other ISR-inducible proteins (Figure 5). It is likely that IFN-γ induces a mild ISR that decreases the ternary complex concentration, but is not enough to observably repress bulk translation (Cagnetta et al., 2019). Cells may also employ additional cell responses other than ISR to tackle IFN-γ stress, such as the unfolded protein response (UPR). The UPR not only converges with the ISR through p-eIF2α, but also induces ISR-independent arms, whereby the transcription factors ATF6 and the spliced form of X-box binding protein 1 (XBP1s) increase expression of genes that reinstate ER homeostasis (Costa-Mattioli and Walter, 2020). Activation of ATF6 but not XBP1s in oligodendrocytes, was reported (Hussien et al., 2015; Mháille et al., 2008; Stone and Lin, 2015). ATF6 deficiency was shown to exacerbates ER stress-induced oligodendrocyte death in the developing CNS of IFN-γ-expressing mice (Stone et al., 2018). The role of ATF6 in oligodendrocyte protection under IFN-γ-stress requires further investigation.

Our data shows that BZA combined with Sephin1 exhibited a faster recovery by increasing remyelinated axons and myelin thickness than the single treatment of BZA. This directed our focus to how Sephin1 promotes remyelination in an inflammatory environment. Sephin1 had little impact on OPC proliferation and differentiation. Therefore, we posit that Sephin1 provides protection to surviving and newly formed oligodendrocytes against inflammation. Although the exact mechanism of how Sephin1 enhances p-eIF2α is controversial (Carrara et al., 2017; Crespillo-Casado et al., 2018), our findings indicate that it enhances the ISR response by increasing p-eIF2α levels in remyelinating oligodendrocytes in the presence of an inflammatory environment. Most importantly, our study uncovers the downstream pathways following ISR enhancement by Sephin1 in oligodendrocytes. We show that Sephin1 elevated the IFN-γ induced p-eIF2α levels, resulting in a reduction of overall protein synthesis, which is protective (Pavitt and Ron, 2012).

Our observation of RNA stress granule formation in IFN-γ stressed cells when treated with Sephin1, helps shed light on the mechanism of Sephin1’s cytoprotective effect. It is widely believed that stress granules are sites of mRNA sequestering during cellular stress (Anderson and Kedersha, 2008; Kedersha and Anderson, 2002). Although stress granules do not directly repress global translation (Khong et al., 2017; Mateju et al., 2020), emerging evidence indicates that stress granules promote cellular survival in diverse disease states by protecting mRNA and proteins from degradation, minimizing energy expenditure, acting in translational quality control, and sequestering proapoptotic factors (Glauninger et al., 2022). Our findings along with previous evidence, raise the possibility that Sephin1’s protective effect on oligodendrocyte is correlated with RNA stress granule formation in response to phosphorylation of eIF2α. Notably, 2BAct, a novel CNS penetrant eIF2B activator (Wong et al., 2019), partially blocked the beneficial effect of Sephin1 on EAE, further supporting our proposed protective mechanism of Sephin1 *in vivo*.

In summary, we observed a significant recovery benefit from the combined treatment of Sephin1 and BZA on inflammatory remyelination and unveiled distinct mechanisms of Sephin1 and BZA on oligodendrocyte lineage cells that underscore this additive effect. These findings have important implications for promoting the ensheathment of demyelinated axons and restoring neuronal function, especially in an inflammatory environment, thereby offering the potential to delay, prevent, or reverse the neurological pathologies of MS. These results should be informative as neuroprotective and neuroreparative therapeutic strategies are being developed for MS and other CNS inflammatory demyelinating disorders.

## MATERIALS AND METHODS

### Animal and drug treatment

Animal use and drug administration were described in our previous study (Chen et al., 2021). GFAP-tTA mice on a C57BL/6J background were mated with TRE-IFN-γ mice on a C57BL/6J background to produce GFAP-tTA; TRE-IFN-γ double-transgenic mice. These mice were given 200 ppm Dox (Envigo, Madison, WI) from conception to prevent transcriptional activation of IFN-γ. The Dox diet was discontinued and replaced with a 0.2% cuprizone diet (Envigo, Madison, WI) starting at six weeks of age. Cuprizone feeding lasted five weeks and then the mice were placed back on normal chow for up to two weeks to allow early remyelination to occur. Concurrently, 8 mg/kg of Sephin1 (i.p.) (#SM1356, Sigma, St. Louis, MO) or 10 mg/kg of BZA (gavage) (#PZ0018, Sigma) were given daily to the GFAP-tTA; TRE-IFN-γ mice, starting from three weeks of cuprizone exposure. The corpus callosum of each mouse was collected two weeks after cuprizone feeding cessation.

EAE was induced in 9-week-old female C57BL/6J mice (Jackson Laboratories, Bar Harbor, ME) by subcutaneous flank administration of 200 μg MOG35-55 peptide emulsified with complete Freund’s adjuvant (CFA) and killed Mycobacterium tuberculosis H37Ra (#EK2100 kit, Hookes Laboratory, Lawrence, MA). Intraperitoneal (i.p.) injections of 200 ng pertussis toxin were given immediately after administration of the MOG emulsion and again 24 hours later. Mice were blindly scored daily for clinical signs of EAE from 0 (no symptoms) to 5 (most severe). Mice were put on a 2BAct diet, which was generously provided by Dr. Carmela Sidrauski’s group (Calico Life Sciences, South San Francisco, CA), two days before EAE induction. Daily treatment with i.p Sephin1 at 8mg/kg were started seven days after EAE induction. The treatment of the animals used in this study was conducted in accordance with the ARRIVE guidelines and in complete compliance with the Animal Care and Use Committee guidelines of Northwestern University.

### OPC isolation and cell culture

Oligodendrocyte precursor cells (OPCs) were isolated and purified following a previously described method (Dugas and Emery, 2013). Briefly, mice brain cortices from 6-7 days old pups were collected, diced, and digested with papain (# 9001734, Worthington, Lakewood, NJ) at 37°C. Cells were triturated, suspended, and sequentially immunopanned at room temperature on plates coated with Ran-2, GalC, and O4 antibodies from a hybridomal supernatant. OPCs sticking to the O4 coated plate were trypsinized and seeded on poly-D-Lysine (pDL)-coated plates in growth media. To get sufficient amount OPCs, the cells were split and plated twice in growth media to proliferate. NIH 3T3 fibroblast cell line was purchased from ATCC (# CRL-1658) and cultured in DMEM with 10% bovine calf serum and 1% penicillin-streptomycin.

### Puromycin incorporation assays

After OPCs were incubated in differentiation media overnight, the cells were treated 200 U/ml of IFN-γ (#485-MI-100, R&D, Minneapolis, MN), and/or 200 nM of thapsigargin (TG, #67526958, Sigma) with or without Sephin1 (50 μM, Sigma) as designated regimens. OPCs were seeded on PDL-coated plates in growth media. One day later, the media were changed to differentiation media and the cells were incubated at 37°C overnight. On the next day, cells were either untreated or treated with IFN-γ or TG, as described above. For puromycin labeling, puromycin (10 μg/ml, Sigma) were added during the last 30 min before harvest.

### Western blot

OPCs were washed twice with PBS and lysed in ice-cold RIPA buffer (Sigma) containing Halt Protease and Phosphatase Inhibitor Cocktail (Thermo Fisher Scientific, PI78441). Lysates were then centrifuged at 12,000 rpm for 20 min at 4°C. A total 20 μg of protein lysates was separated by 4-12% SDS-PAGE (Bio-Rad, 4561095) and transferred to a nitrocellulose membrane. The following primary antibodies were used: anti-p-eIF2α (Abcam, ab32157, 1:2000), anti-T-eIF2α (Cell Signaling, 9722s, 1:1000), anti-puromycin (Millipore, MABE343, 1:2000), anti-BIP (Cell Signaling, 3177s, 1:1000), anti-GADD34 (Proteintech, 10449-1-AP, 1:500), anti-ATF4 (Santa Cruz, sc-390063, 1:500), anti-CHOP (Thermo Fisher, MAI-250, 1:500), anti-XBP-1-spliced (Cell Signaling, 82914s, 1:1000), and anti-actin (Sigma, A2066, 1:2000). Quantification of Western blot bands were performed by Image Lab Software (Bio-Rad).

### Cell treatments and Immunocytochemistry

OPCs were grown on PDL-coated 4-well glass culture slides (BD, 354104). To study OPC differentiation, we exchanged medium to one without differentiation factors (T3) and exposed cells to 200 U/ml IFN-γ with the treatment of 500 nm BZA (#PZ0018, Sigma), and/or 50 uM Sephin1 (a gift from InFlectis Biosciences) for 24, 48 and 72 hrs. To induce cellular stress, cells were treated for one hour with 500 μM sodium arsenite (#S7400, Sigma), 200 nM TG, or 200 U/ml IFN-γ. A longer treatment was conducted for 8, 16, 24, or 48 hours with 200 U/ml IFN-γ, and/or 50 uM Sephin1.

Cells were washed twice with PBS and fixed with ice-cold 4% PFA for 10 min at RT. Following fixation, cells were washed three times with PBS for 5 min, and then blocked in PBS supplemented with 5% BSA, 1% normal donkey serum (DNS), and 0.1% Triton X-100 for one hour. Primary antibodies were diluted in block solution and incubated at 4°C overnight. The following antibodies were used: Rabbit anti-G3BP1 (Abcam, ab181150, 1:500) and mouse anti-O1 (R&D, MAB1327, 1:500), anti-PDGFR-alpha (BD Biosciences, 558774, 1:100), anti-MBP (Abcam, ab24567, 1:700). Following primary antibody incubation, the cells were washed with PBS and probed with Alexa Fluor-conjugated secondary antibodies (Thermo Fisher Scientific,1:500) for 1hr at RT. Cells were then washed with PBS and mounted using HardSet™ Antifade Mounting Medium with DAPI (VECTASHIELD, H-1500-10). Cells were viewed on a Zeiss LSM 880 confocal microscope.

### Histology (immunofluorescent staining and electron microscopy) and imaging

The histology studies were previously described (Chen et al., 2021). In short, cerebellums were harvested first and then the same mice were the perfused with 4% paraformaldehyde (PFA, Electron Microscopy Sciences, Hatfield, PA) in PBS for 15 min. The brains were removed and cut coronally at approximately 1.3 mm before the bregma. The posterior parts of the brains were postfixed with 4% PFA overnight, embedded in an OCT compound and cryosectioned in a series of 10 μm. The immunostaining protocol is described in Chen et al., 2021. Primary antibodies included the following: anti-MBP (Abcam, ab24567, 1:700), anti-ASPA (Genetex, GTX113389, 1:500), anti-Ki67 (Abcam, AB15580, 1:100), anti-PDGFR-alpha (BD Biosciences, 558774, 1:100), anti-cc1 (Calbiochem, OP80, 1:200) and anti-p-eIF2α (Abcam, ab32157, 1:500). The fluorescent stained sections were scanned with an Olympus VS-120 slide scanner and quantified by Image J. At least three serial sections of corpus callosum were quantified. The representative fluorescent images were acquired under a Zeiss LSM 880 confocal microscope.

The anterior parts of the brains were immersed in EM buffer for two weeks at 4°C (Chen et al., 2021), then dehydrated in ethanol, cleared in propylene oxide, and embedded in EMBed 812 resin (Electron Microscopy Sciences). Grids were examined on an FEI Tecnai Spirit G2 transmission electron microscope. The total percentage of remyelinated axons, averaged from 10 images (area = 518.3504 μm2) in each mouse, was calculated. G-ratios were calculated as the ratio of the inner diameter to the outer diameter of a myelinated axon; a minimum of 300 fibers per mouse was analyzed.

### Quantitative real-time reverse transcription PCR (RT-qPCR)

The RT-qPCR procedure was described previously (Chen et al., 2021). Total RNA was isolated from the cerebellum. RT-qPCR was performed on a CFX96 RT-PCR detection system (Bio-Rad) using SYBR Green technology. Results were analyzed and presented as the fold-induction relative to the internal control primer for the housekeeping gene GAPDH. The primers (5’-3’) for the mouse gene sequences were as follows: Gapdh-f: TGTGTCCGTCGTGGATCTGA, Gapdh-r: TTGCTGTTGAAGTCGCAGGAG; Ifng-f: GATATCTGGAGGAACTGGCAAAA, Ifng-r: CTTCAAAGAGTCTGAGGTAGAAAGAGATAAT.

### Super-resolution imaging and processing

Structured illumination microscopy (SIM) images were obtained using a Zeiss Elyra 7 equipped with dual pco.edge 4.2 sCMOS cameras. Collected light split using a Long ass 560 filter and 405, 488, and 561nm laser lines were used for excitation. Images were acquired using Zen Black Software (Zeiss). SIM images were processed using dual iterative reconstruction (SIM2, Zeiss) in Zen Black using Standard Live preset parameters.

### Statistical analysis

All results are expressed as mean±SEM. Statistical significance for these data was determined by t-tests, analysis of variance (ANOVA) followed by Student’s test and the Bonferroni method for multiple group experiments. Significance levels were set at p < 0.05.

## ACKNOWLEDGEMENTS

We thank Erdong Liu and Nia Stewert for technical assistance, and we thank Dr. Jeff Twiss for advice on detecting RNA stress granules. We thank Inflectis Bioscience for providing Sephin1. This study was supported by NIH/NINDS R01 NS034939 (BP), the Dr. Miriam and Sheldon G. Adelson Medical Research Foundation (JRC and BP), the Rampy MS Research Foundation (JRC and BP), and National Multiple Sclerosis Society Career Transition Fellowship TA-2008-37043 (YC).

## SUPPLEMENTAL FIGURES

**Figure 1-figure supplement 1.**
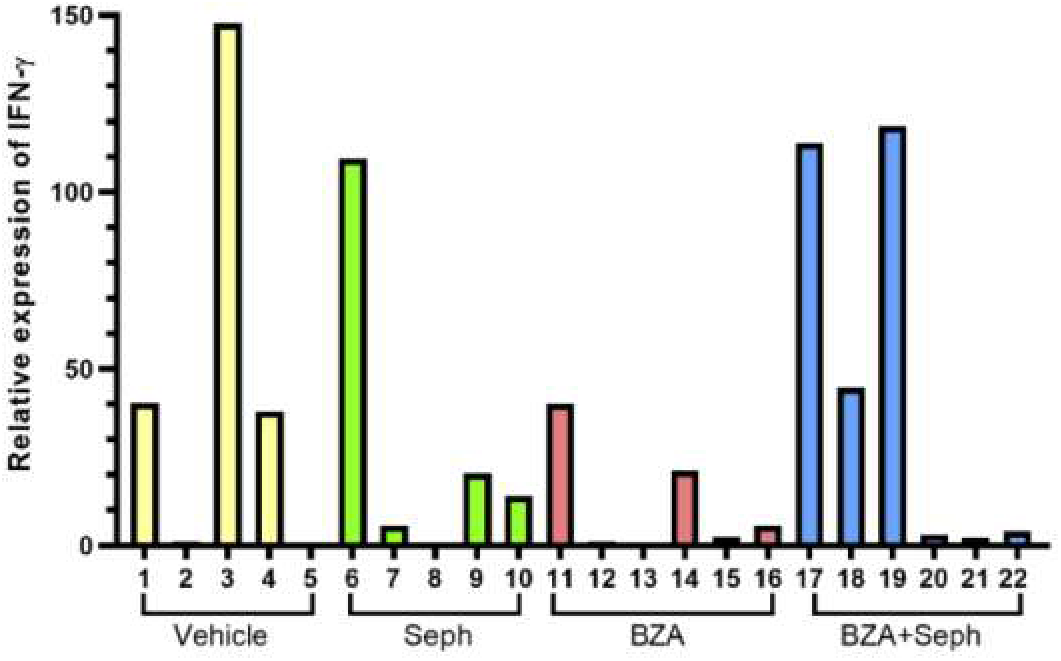
GFAP-tTA;TRE-IFN-γ mice express IFN-γ after release from doxycycline. The expression levels of IFN-γ in the cerebellum two weeks after cuprizone withdrawal in each treatment group (RT-PCR). Each bar is an individual mouse. Mouse #1, 3, 4, 6, 7, 9, 10, 11, 14, 16, 17, 18 and 19 were selected for further examination.

**Figure 3-figure supplement 2.**
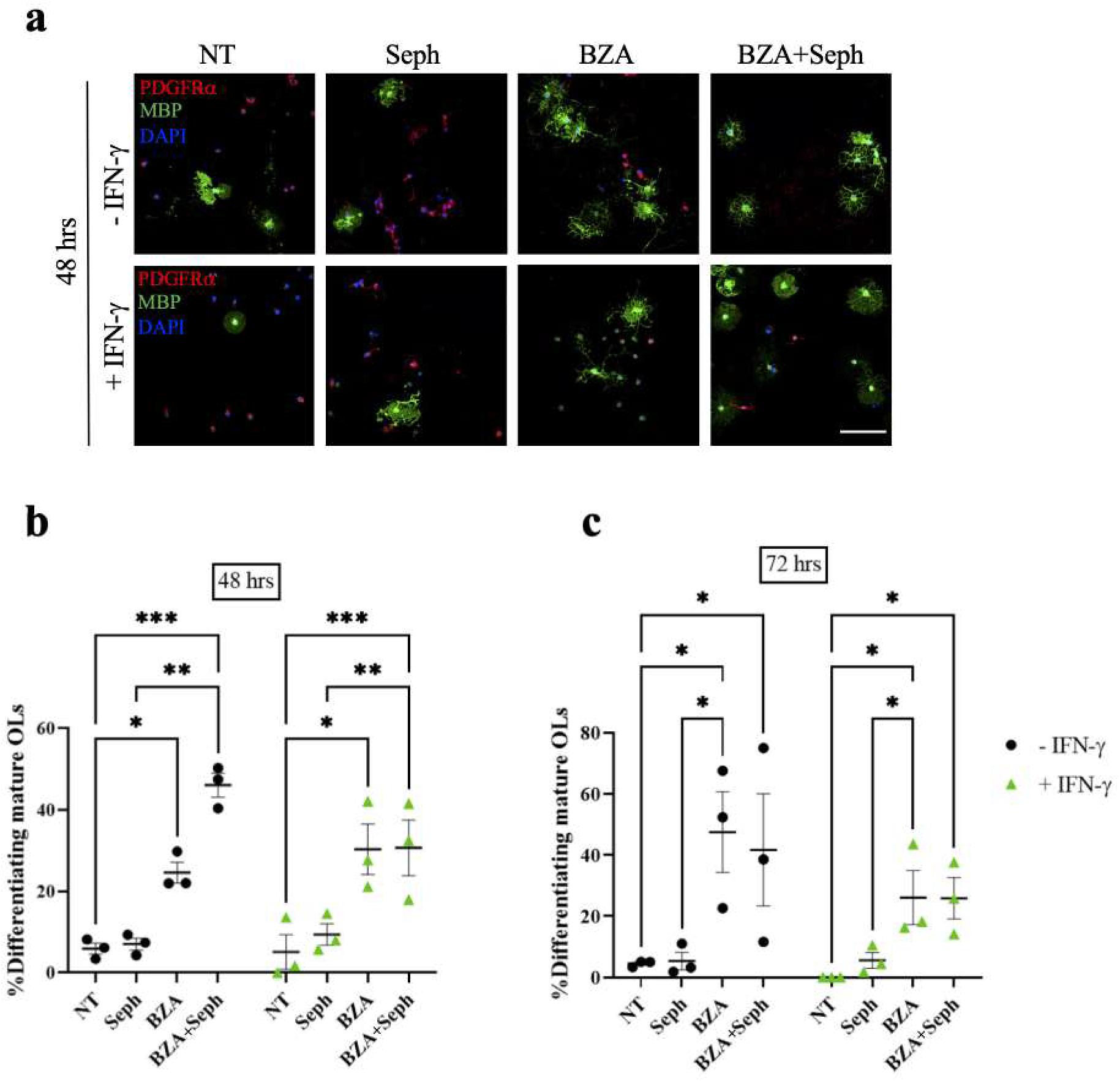
Combined treatment of Sephin1 and BZA promote OPC differentiation at 48 and 72 hours. (**a**) PDGFR-α and MBP immunostaining of OPCs in cultures that were seeded for 48 hours. Cells exposed to IFN-γ (+ IFN-γ) were treated with non-treatment (NT), Sephin1 (Seph1), BZA, or BZA plus Seph1. Cells not exposed to IFN-γ (-IFN-γ) were used as controls. Scale bar: 100μm. Quantification of percentage of cells positive for MBP (differentiating oligodendrocytes) over the total number of oligodendrocyte lineage cells at 48 hours (**b**) and 72 hours (**c**). *p < 0.05, **p < 0.01.

**Figure 5-figure supplement 3.**
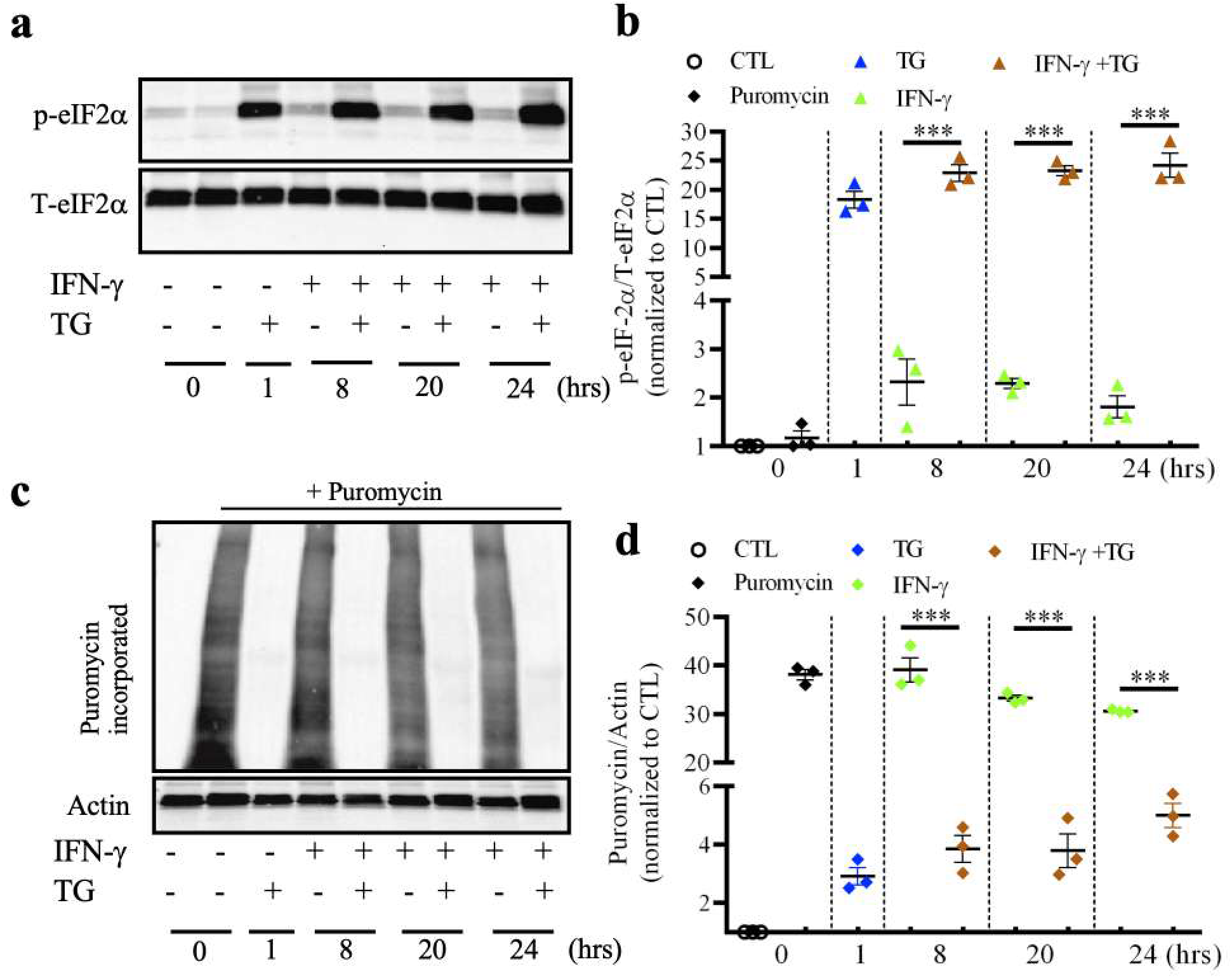
IFN-γ combined with TG is able to induce the ISR and reduces the overall protein translation in mouse oligodendrocytes. **(a)** Western blot of oligodendrocytes treated by IFN-γ and/or TG. **(b)** Quantification of a Western blot for p-eIF2α levels. **(c)** Western blot of puromycin incorporation assay using an anti-puromycin antibody. OPCs were differentiated in differentiation media overnight and then treated with IFN-γ and/or TG. Puromycin was added 30 min before harvesting for puromycin labeling. **(d)** Quantification of Western blot. Data are mean ± SEM from three biological isolations and technique replicates. ***p < 0.001. **Figure 5-figure supplement 3-source data 1**. Western-blots images of p-eIF2α and puriomycin in oligodendrocytes exposed to IFN-γ and/or Thapsigargin (TG).

**Figure 6-figure supplement 4.**
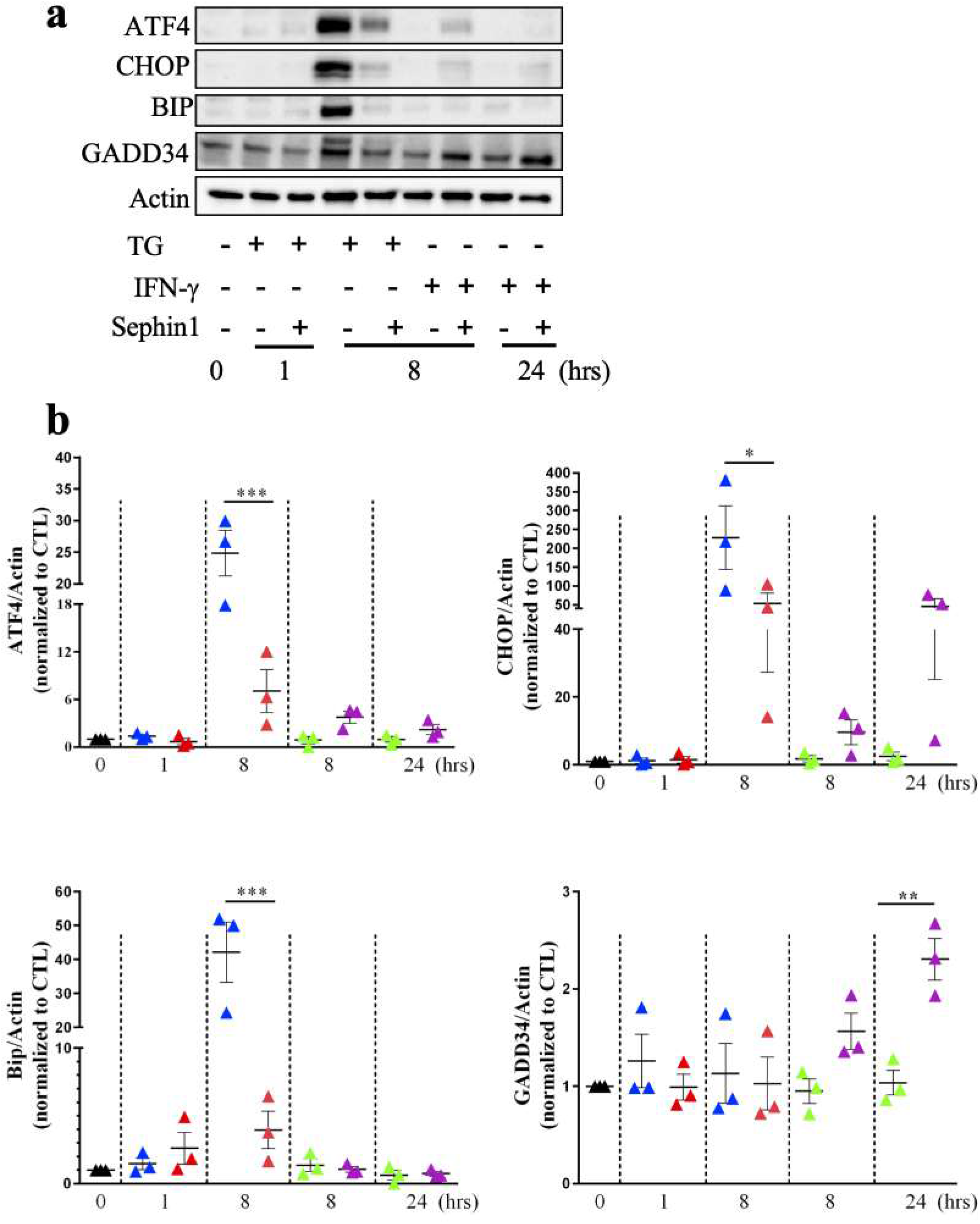
Sephin1 increases the expression of ISR downstream targets. Immunoblot analysis of p-eIF2α, total eIF2α and ISR response (ATF4, BIP, GADD34, and CHOP) in either TG or IFN-γ exposed developing oligodendrocytes. Cells were treated with Sephin1 for 1, 8 and 20 hours. **Figure 6-figure supplement 4-source data 1**. Western-blots images of downstream ISR proteins in oligodendrocytes exposed to thapsigargin (TG) or IFN-γ with or without Sephin1 treatment.

**Figure 7-figure supplement 5.**
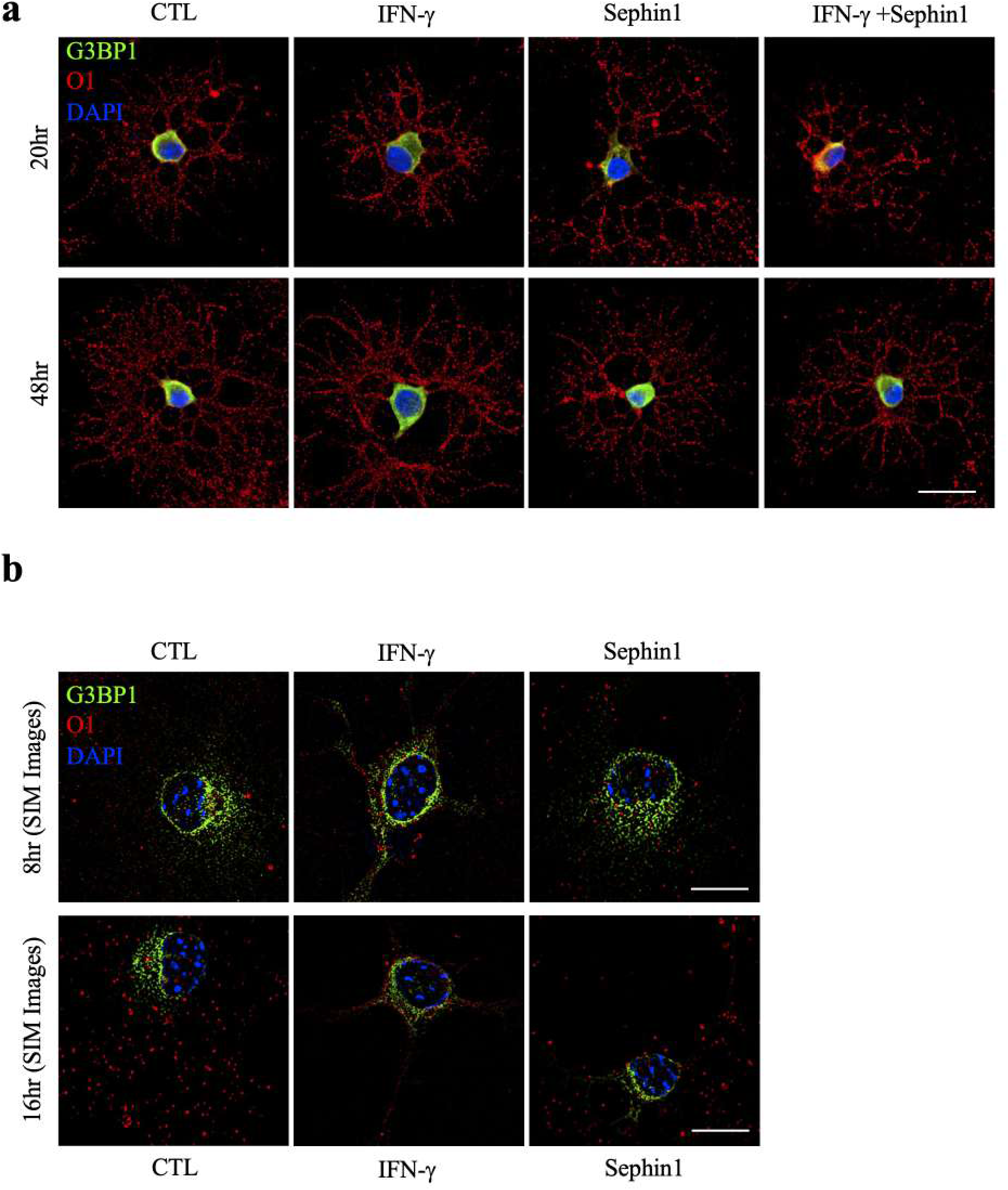
Stress granules are not observed in IFN-γ exposed oligodendrocytes when treated with Sephin1 after 20 and 48 hours. (**a**) Representative confocal images of the stress granule marker G3BP1 (green) and oligodendrocyte marker O1 (red) in purified oligodendrocytes treated for 1hr with IFN-γ and/or Sephin1 for 20 and 48 hours. Scale bar indicates 20 μm. (**b**) Representative SIM images of G3BP1 (green) and O1 (red) in purified oligodendrocytes with a single treatment at 8 and 16 hours. Nuclei are stained with DAPI (blue). Scale bar indicates 10 μm.

